# POFUT2-mediated *O*-glycosylation of MIC2 is dispensable for *Toxoplasma gondii* tachyzoites

**DOI:** 10.1101/391623

**Authors:** Sachin Khurana, Michael J. Coffey, Alan John, Alessandro D. Uboldi, My-Hang Huynh, Rebecca J. Stewart, Vern B. Carruthers, Christopher J. Tonkin, Ethan D. Goddard-Borger, Nichollas E. Scott

**Author notes:** To whom correspondence and requests for materials should be addressed E.D.G.-B., C.J.T or N.E.S. these authors contributed equally to the work (Sachin Khurana, Michael J. Coffey).

## Abstract

*Toxoplasma gondii* is a ubiquitous obligate intracellular eukaryotic parasite that causes congenital birth defects, disease of the immunocompromised and blindness. Protein glycosylation plays an important role in the infectivity and evasion of immune response of many eukaryotic parasites and is also of great relevance to vaccine design. Here, we demonstrate that MIC2, the motility-associated adhesin of *T. gondii*, has highly glycosylated thrombospondin repeat domains (TSR). At least seven *C-*linked and three *O*-linked glycosylation sites exist within MIC2, with >95% occupancy at *O*-glycosylation sites. We demonstrate that the addition of *O*-glycans to MIC2 is mediated by a protein *O*-fucosyltransferase 2 homologue (TgPOFUT2) encoded by TGGT1_273550. While POFUT2 homologues are important for stabilizing motility associated adhesins and host infection in other apicomplexan parasites, in *T. gondii* loss of TgPOFUT2 has only a modest impact on MIC2 levels and the wider proteome. Consistent with this, both plaque formation and tachyzoite infectivity are broadly similar in the presence or absence of TgPOFUT2. These findings demonstrate that TgPOFUT2 *O*-glycosylates MIC2 and that this glycan is dispensable in *T. gondii* tachyzoites.

## Introduction

The phylum Apicomplexa is comprised of a large group of obligate intracellular eukaryotic parasites, many of which have medical and agricultural significance. Pathogenic apicomplexan species include; *Plasmodium* spp. (malaria), *Cryptosporidum* spp. (cryptosporidosis), *Theileria* spp. (theleriosis), *Basbesia* spp. (babesiosis) and, the most ubiquitous of all, *Toxoplasma gondii* (toxoplasmosis). *T. gondii* infects 30-80% of the human population and, whilst largely self-limiting in healthy individuals, it can cause major problems in the immunosuppressed and lead to congenital birth defects if contracted whilst pregnant (1). Some countries have extremely high rates of progressive blindness caused by toxoplasmic retinopathy, which have no curative treatment (1–3).

All apicomplexan parasites must migrate through host tissues and invade cells to survive and to cause disease. Central to this process is a unique form of cellular locomotion termed ‘gliding motility’. Gliding motility requires the apical release of transmembrane adhesins from microneme organelles onto the parasite surface (4), which then provides anchor to the extracellular environment and/or host cells. The current model posits that motility is initiated when an actomyosin-based ‘glideosome’, which lies just underneath the plasma membrane, binds to the cytoplasmic tails of adhesins and drags them to the rear of the parasite, exerting a forward-acting force thus driving forward motion (5–7).

Motility-associated adhesins in Apicomplexa vary between species and differ in expression across the various life-cycle stages, reflecting the diversity of host cells that are targeted by these parasites. For example, *Plasmodium* spp. use a range of adhesins that specifically bind to erythrocyte receptors in asexual blood stages and other adhesins in stages that infect mosquitoes or the human liver (8,9). While little is known about the host cell receptors of *T. gondii*, a diverse array of putative adhesins are contained within their micronemes, which goes some way to explaining the remarkably diverse range of host species and cell types that can be infected by this zoonotic parasite. Common features exist among these motility and invasion-associated adhesin proteins, including the recurrence of one or more thrombospondin repeat (TSR) domains (10).

Apicomplexan TSR-containing adhesin proteins are the only examples of non-metazoan TSR domains. Metazoan TSRs are glycosylated in the ER with the *O*-linked glycan α-D-glucopyranosyl-1,3-β-L-fucopyranoside (GlcFuc) and *C-*linked α-D-mannopyranosides (*C-*Man) (11). Protein *O*-fucosyltransferase 2 (POFUT2) initiates metazoan TSR *O*-glycosylation and plays an important role in the folding and stabilization of these proteins (12–16). Recently, TSR *O*-glycosylation has also been observed in *Plasmodium falciparum* and *Plasmodium vivax* (17,18): this *O*-glycan is initiated by a homologue of POFUT2 and is important for stabilizing proteins with TSR domains and efficient host infection (19). *O*-Glycosylation of TSR domains may also be an important consideration in vaccine design (20). Endogenous TSR *O*-glycosylation has yet to be observed in *T. gondii*, however it does encode a putative POFUT2 (TGGT1_273550) and the recombinant expression of *T. gondii* proteins in CHO cells does results in modification of parasite protein with the GlcFuc and *C-*Man glycans (21). Here, we generate a *T. gondii* line to facilitate bulk purification of micronemal protein 2 (MIC2) directly from tachyzoites in order to map endogenous glycosylation sites on this motility-associated adhesin that possesses six TSR domains. We go on to demonstrate that a *T. gondii* POFUT2 is encoded by TGGT1_273550 and that loss of *O*-glycosylation has only a modest impact on MIC2 levels, the wider tachyzoite proteome and infectivity.

## Results

### MIC2 is glycosylated at multiple sites to high occupancy

To explore the glycosylation of MIC2 we generated a *T. gondii* line in which the MIC2 associated protein (M2AP) possessed a C-terminal strep-FLAG tandem affinity purification (SF-TAP) tag to allow for convenient purification of the M2AP-MIC2 complex. Western blot using FLAG antibodies confirmed that M2AP was tagged (Figure 1A). Further, we showed using immunofluorescence that M2AP-SF-TAP largely co-localizes with MIC2, as expected (Figure 1B). The SF-TAP tagged M2AP enabled simple high-yielding batch purification of the M2AP-MIC2 complex from bulk tachyzoite cultures (Figure 1C). We characterized MIC2 glycosylation (Figure 2A) using a combination of enzymatic digestions (trypsin, gluC, and sequential gluC then trypsin) together with multiple fragmentation approaches, including higher-energy collision dissociation and electron-transfer and higher-energy collision dissociation fragmentation approaches (HCD and EThcD (22) respectively, Supplementary Table 1). Using this approach, we detected multiple *C-*glycosylation and *O*-glycosylation events in five of the six TSR domains of MIC2 (Figure 2A and B, Supplementary Figure S1). Within these TSR domains a total of five previously unreported *C-* glycosylation events were observed (Figure 2B) corresponding to W^276^, W^279^, W^348,^ W^351^ and W^479^ with all except W^479^ lying within the *C-* mannosyltransferase consensus motifs (WXXW/C (23)). Electron transfer-based fragmentation was used to localize the labile *O*-glycosylation events to three sites within MIC2: S^285^ (Figure 3A), S^485^ (Figure 3B), and T^546^ (Figure 3C and D). We compared the abundance of unmodified and modified peptides covering these sites to better understand *O*-glycan occupancy (Supplementary Table 2) and all of the *O*-glycosylated peptides were found to be the dominant observable species (Figure 3E, F and G). To surmise, MIC2 is endogenously modified with at least three *O*-glycans (GlcFuc) and seven *C-*glycans (*C-*Man) (Figure 2 and 3) and that the *O*-glycan occupancy is greater than 95%.

**Figure 1:**
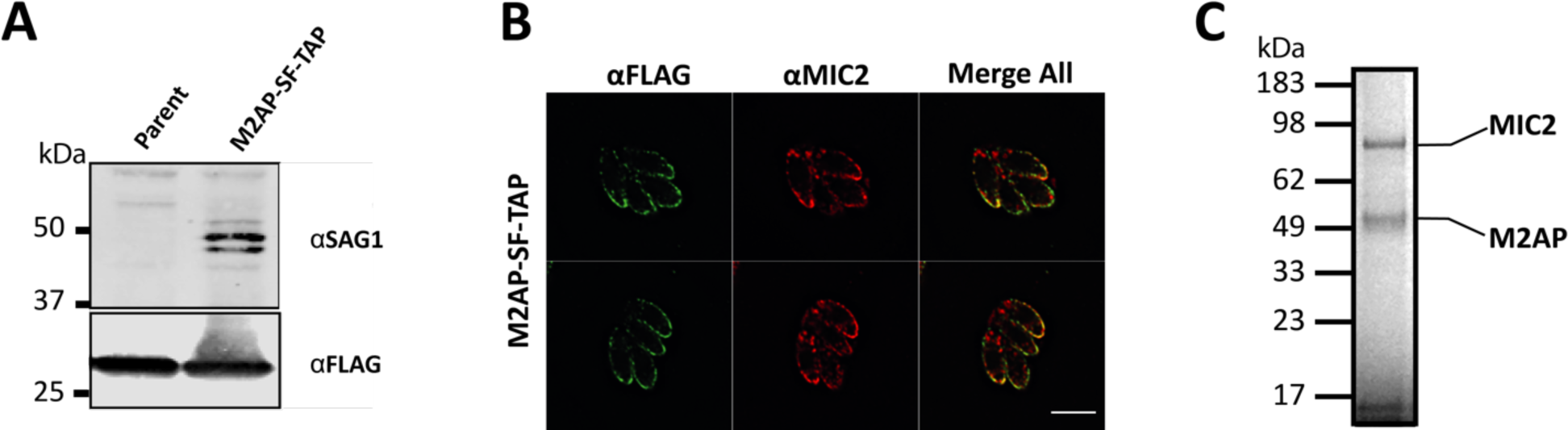
Establishment and validation of M2AP SF-TAP *T. gondii* line. A) The addition of the SF-TAP tag enables the detection M2AP within the *T. gondii* line M2AP SF-TAP compared to the parental line. B) M2AP, detected using *α*-Flag, co-localises with MIC2 within tachyzoites. C) Enrichment of M2AP SF-TAP tagged protein enables the isolation of the MIC2-M2AP complex to high purity as determined by Coomassie stained gels.

**Figure 2:**
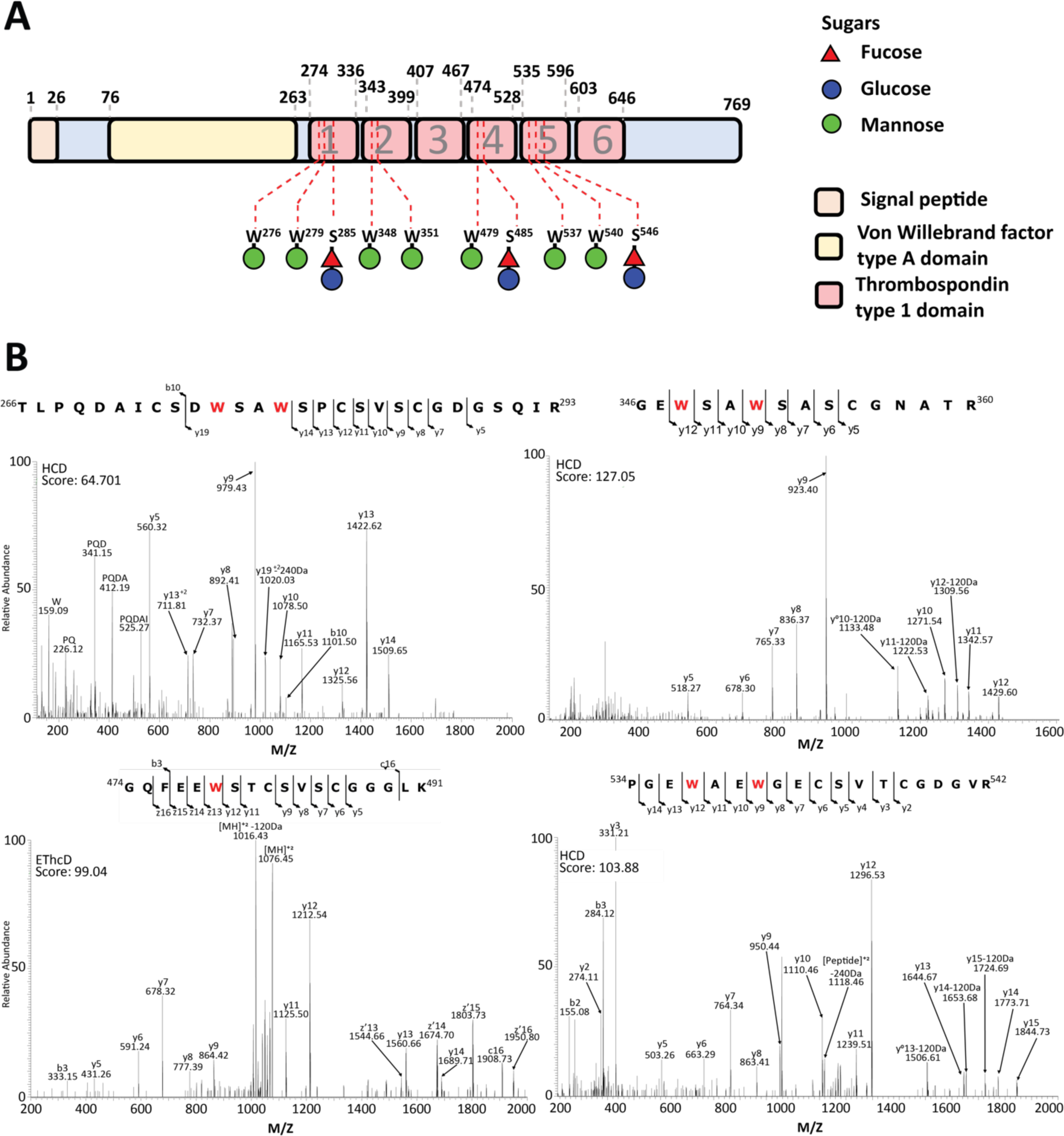
Characterisation of MIC2 glycosylation events. A) Within MIC2 ten glycosylation sites were identified corresponding to 7 *C-*glycosylation events and 3 GluFuc sites of modification. The residues modified and the corresponding carbohydrate modification are mapped to the MIC2 sequence. All observed glycosylation events all lie within the TSR domains. B) Peptides containing observed the *C-*glycosylation events W^276^, W^279^, W^348,^ W^351^, W^479^, W^537^ and W^540^ are shown.

**Figure 3:**
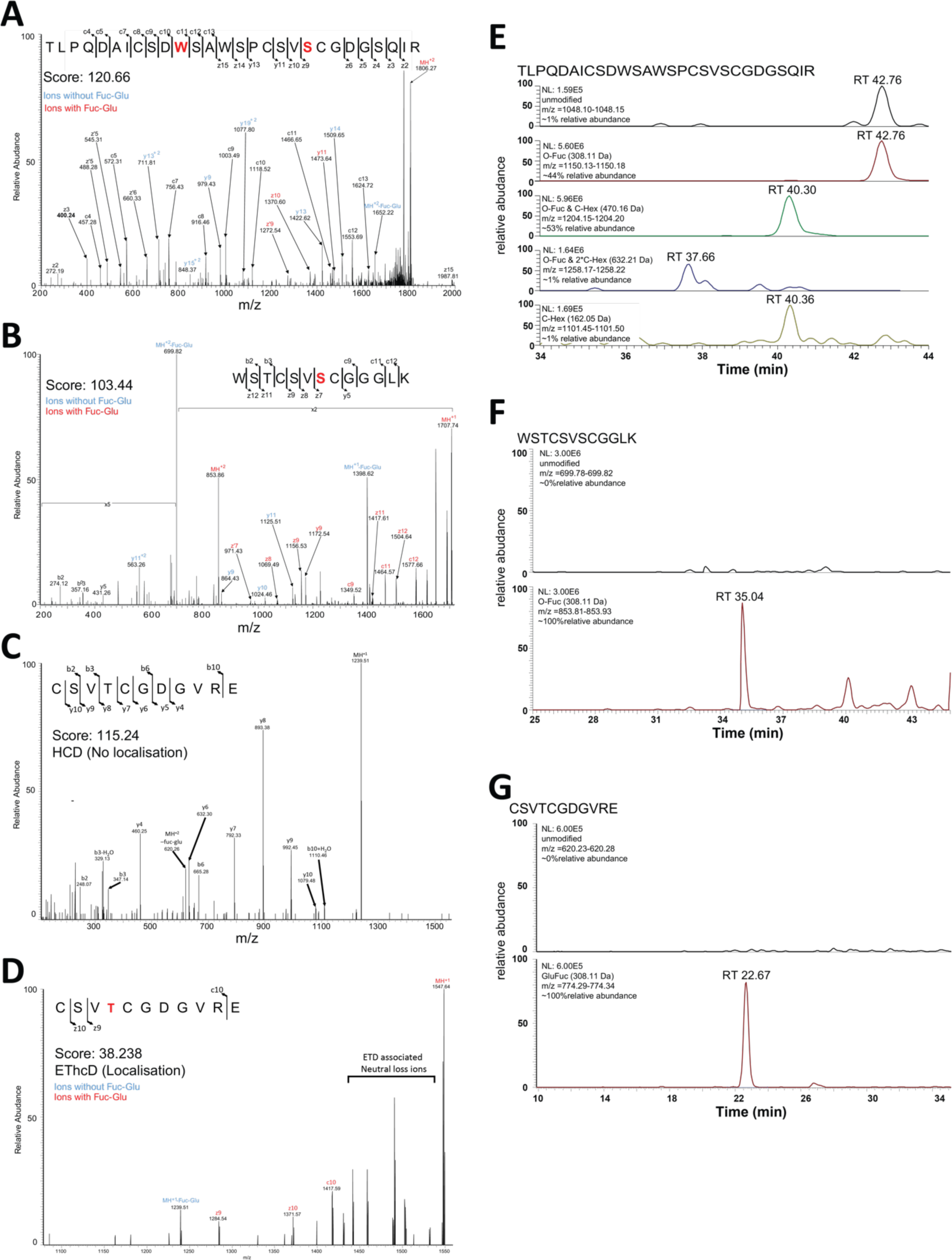
Site of *O-*fucosylations of MIC2 are modified at a high occupancy. A) Analysis of tryptic digested MIC2 with EThcD fragmentation enabled identification of the C-glycosylated and GluFuc within ^266^TLPQDAICSDWSAWSPCSV<u>S</u>CGDGSQIR^293^, with the site of GluFuc localised to S^285^ B) Analysis of trypsin and GluC digested MIC2 with EThcD fragmentation enabled identification of glycosylated ^479^WSTCSV<u>S</u>CGGGLK^491^, with the site of GluFuc localised to S^485^. Analysis of GluC digested MIC2 using a combination of HCD (C) and EThcD (D) fragmentation enabled the localisation of GluFuc to T^546^ on the glycopeptide ^543^CSV<u>T</u>CGDGVRE^553^. The GluFuc containing glycoforms of these peptides were the most abundant forms observed (E, F and G) supporting these sites are occupied at high occupancy.

### TGGT1_273550 encodes the *T. gondii* POFUT2

We were interested in determining the enzyme responsible for deposition of *O*-glycans on MIC2. A BLAST search of the *T. gondii* GT1 genome using *Homo sapiens* POFUT2 (CAC24557.1, [28]) as the query sequence suggested that the protein encoded by TGGT1_273550 was most likely the *T. gondii* POFUT2 homologue. To investigate the function of TgPOFUT2 we first introduced a C-terminal triple HA (HA_3_) epitope tag using single crossover recombination (Figure 4A). This allowed us to monitor disruption of TgPOFUT2 using CRISPR/Cas9-based gene knockout (Figure 4A and Supplementary Figure S2A-B), which was all performed in the M2AP-SF-TAP genetic background so as to provide a convenient means to purify and analyze MIC2 in the absence of TgPOFUT2(*Δtgpofut*). We first looked to see if loss of TgPOFUT2 impacted MIC2 protein levels, as was observed for TRAP in *P. falciparum* (19). Interestingly, we also observed reproducible faster migration of MIC2 by SDS-PAGE, perhaps reflecting the small mass change and/or an increase in polypeptide hydrophobicity that results from loss of multiple *O*-glycans (Figure 4Bi) and found a small, yet consistent, reduction in abundance as compared to parental lines (Figure 4Bii). We then used the SF-TAP handle to purify the MIC2-M2AP complex from both parental and *Δtgpofut2* lines as before (Figure 4C), subjecting these samples to trypsin digestion followed by MS analysis and in doing so could observe no difference in the ability of MIC2 to precipitate M2AP suggesting no difference in interaction between these two proteins. Monitoring the MIC2 glycopeptide^266^TLPQDAICSD<u>W</u>SA<u>W</u>SPCSV<u>S</u>CGDGSQIR^293^, which can be both *C*- and *O*-glycosylated, provided insights into glycosylation changes within the *Δtgpofut* line (Figure 4D panels 2 to 4). We could not observe any GlcFuc containing forms of this peptide, yet we were readily able to identify the unmodified form of this peptide (retention time 53.06 min), and both the singly and doubly *C*-glycosylated forms of this peptide (retention time 49.58 and 45.25 min, respectively) (Figure 4D panel 5 and 6, Supplementary table 3 and 4). Collectively, this data suggests that the *T. gondii* POFUT2 is encoded by TGGT1_273550 and that TSR *O*-glycosylation plays only a minor role in regulating MIC2 abundance.

**Figure 4:**
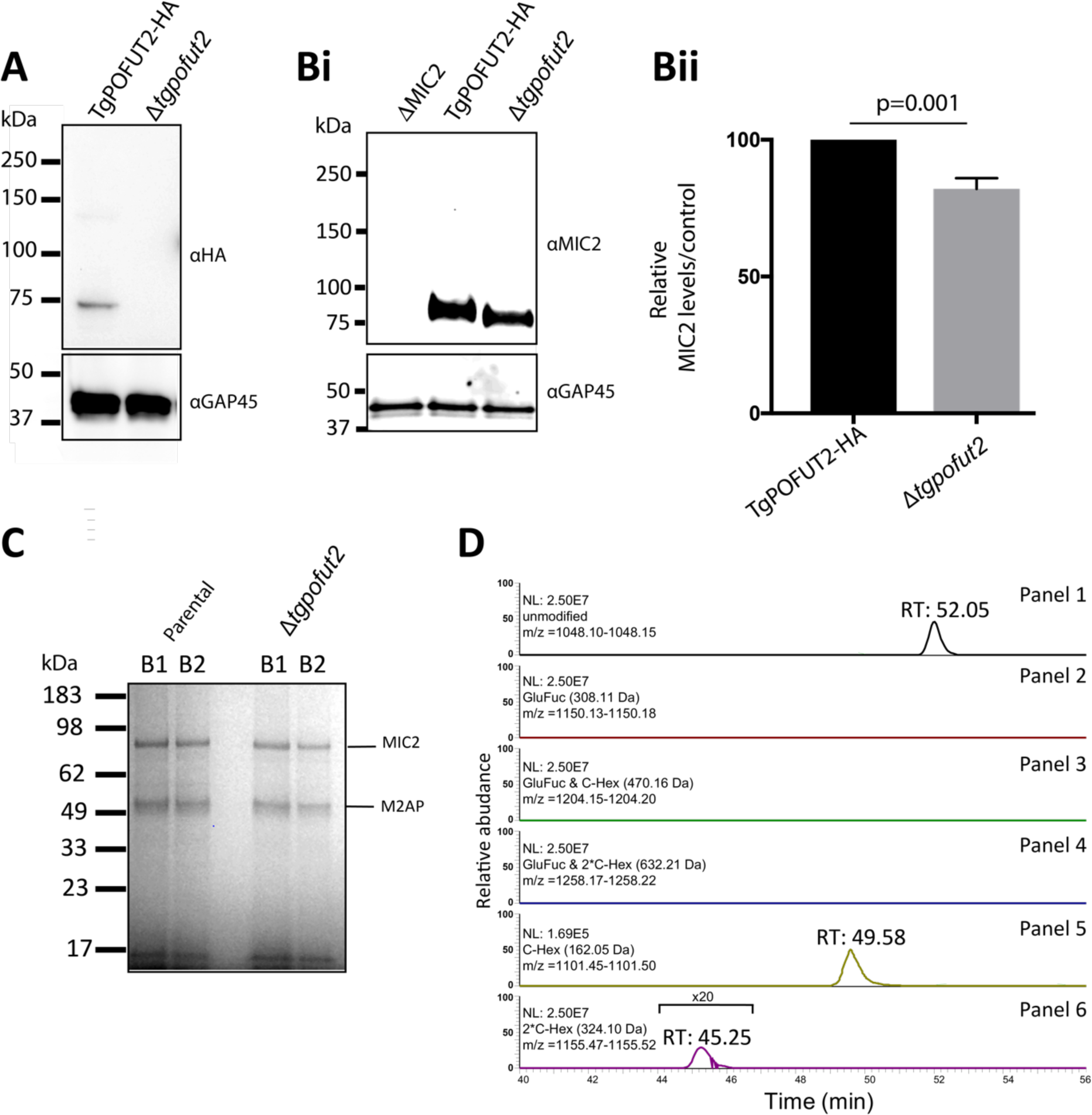
TGGT1_273550 encodes the *T. gondii* POFUT2. A) Endogenously tagging tgPOFUT2 enabled the detection of tgPOFUT2 confirming the expressing in tachyzoites. This band is absent within a *Δtgpofut2* line. B) Loss of POFUT2 leads to a change in the migration of MIC2 and a small, ~20%, yet statically significant change in the amount of MIC2. C) Isolation of M2AP enables the co-isolation of MIC2 within *Δtgpofut2* line at comparable levels, B1 and B2 corresponding to two independent isolations from biological replicates. D) For the peptide ^266^TLPQDAICSDWSAWSPCSV<u>S</u>CGDGSQIR^293^, unmodified, single and doubled C-glycosylated peptides were identified yet no GluFuc containing species were observed. Extracted ion chromatograms demonstrates these identified forms are clearly detectible in panel 1, 5 and 6 while even at the MS1 level no ions corresponding to the GluFuc glycoforms were observed in panel 2 to 4.

### Disruption of TgPOFUT2 results in modest alterations across the proteome

Since glycosylation systems can target multiple protein substrates, and because this modification is a known stabilizing factor, the loss of TgPOFUT2 might disrupt a range of proteins beyond MIC2 (24,25). To gain a better understanding of underlying changes that result from the loss of *O*-glycosylation in *T. gondii* we conducted a global proteomics analysis comparing the parental strain to *Δtgpofut2* using Label-Free Quantitative (LFQ) proteomics (Supplementary table 5) (26). A total of 3839 unique *T. gondii* proteins were identified across biological replicates with >3000 quantified proteins within each of the five parental and *Δtgpofut2* replicates. We observed only modest changes in the proteome in response to the loss of TgPOFUT2 (Figure 5A). These modest changes are reflected in the Pearson correlation (>0.95) observed between samples (Supplementary Figure S3). Using conventional thresholds (fold change > ±1 fold and P-value >0.05) we observed a total of 26 proteins that decreased in abundance while 4 increased in abundance within *Δtgpofut2* compared to parent (Figure 5A). Even with less stringent thresholds (fold change > ±1 fold and P-value >0.075) few additional alterations are observed across the proteome of the *Δtgpofut2* strain (Figure 5B, 37 decreasing, and 21 increasing respectively). Although modest, these alterations are consistent across biological replicates (Figure 5B) and suggest that loss of TSR *O*-glycosylation leads to small but real changes in the *T. gondii* proteome. Importantly, no TSR containing proteins were observed to be undergo significant changes in abundance in response to the loss of TgPOFUT2.

**Figure 5:**
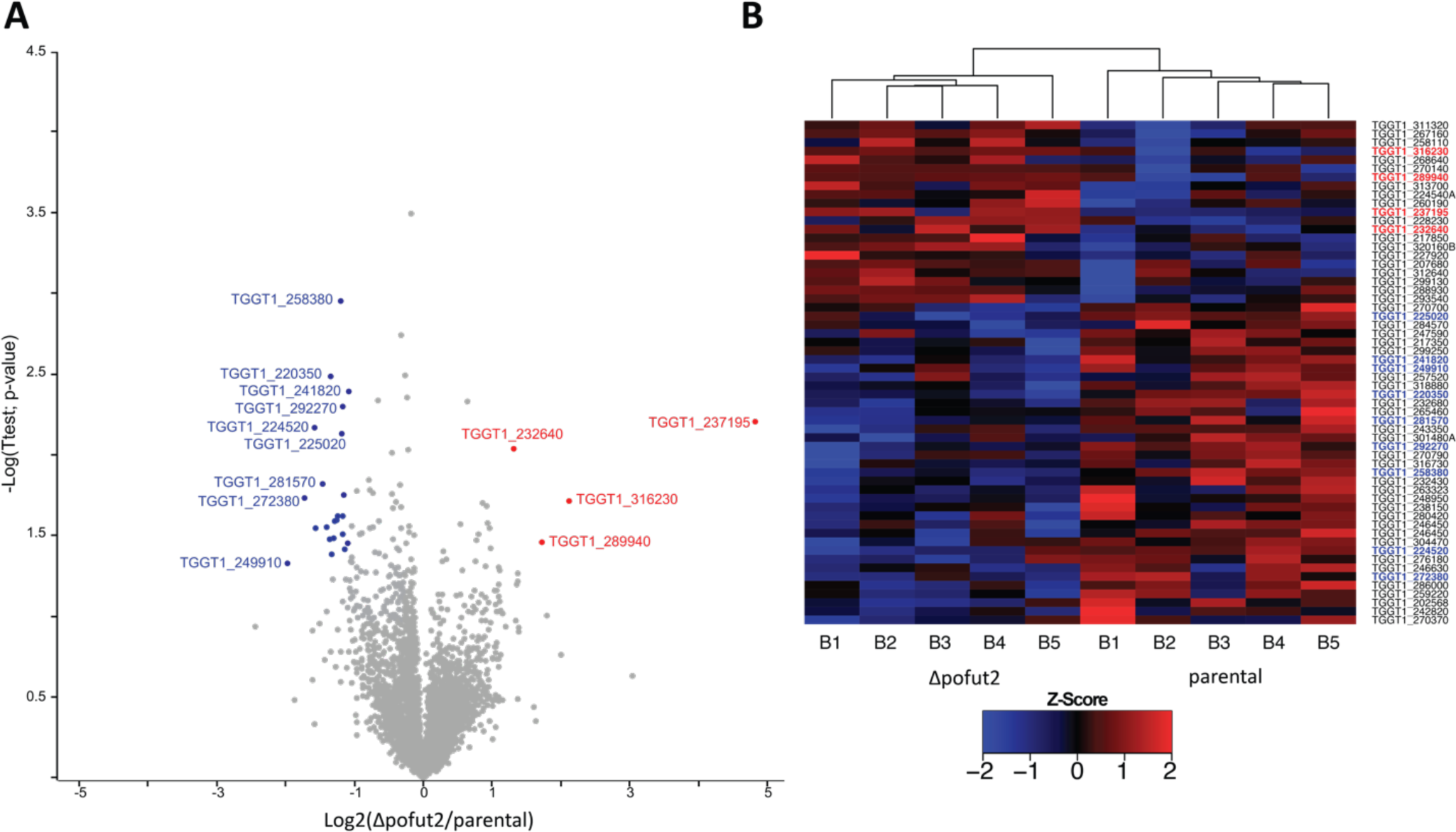
Quantitative proteomic analysis of *Δtgpofut2* compared to the parental line. Label-free quantification of isolated tachyzoites was undertaken to compare *Δtgpofut2* to the parental line. A) Identified proteins are presented as a volcano plot depicting mean label free quantitation (LFQ) intensity ratios of *Δtgpofut2* versus the parental line plotted against logarithmic *t* test *p* values from five biological experiments of each line. B) Heat map of z-scored values of the proteins observed to change between the *Δtgpofut2* and the parental line demonstrating the consistency of these changes across experiments.

### TSR *O-*glycosylation is not important for lytic stage growth, MIC2 trafficking or host cell invasion

We furthered our phenotypic analysis to reveal if TgPOFUT2 contributes to lytic stage growth in relation to what is known about MIC2 biology in *T. gondii*. To do this we first performed plaque assays whereby parental and *Δtgpofut2* parasites were left to grow on human fibroblast (HFF) monolayers over 7 days and then zones of lysis (plaques) quantitated. Here we confirmed, as previously described (10), that parasites lacking MIC2 cause a reduction in plaque size, as compared to parental line (Figure 6A) (27,28). However, *Δtgpofut2* tachyzoite plaques appear normal in morphology and their size is statistically no different than parent (Figure 6Ai and ii), nor does the plaquing capacity (as a percentage of inoculated tachyzoites) change (Figure 6Aiii).

**Figure 6:**
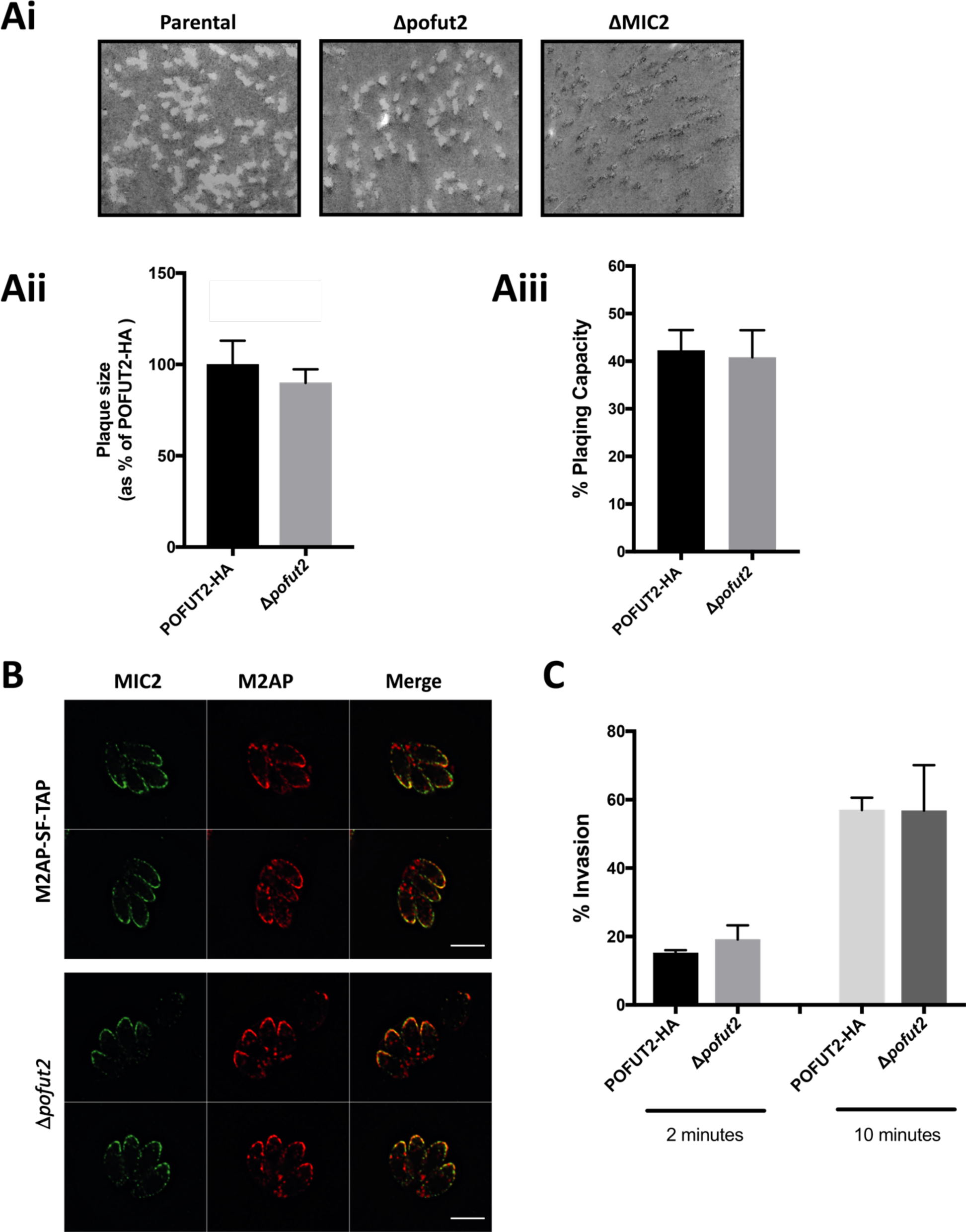
Phenotypic analysis of *Δtgpofut2* compared to the parental line. Ai) Morphological assessment of plaque assays comparing the parental line, *Δtgpofut2* and *ΔMIC2* Aii) Numerical assessment of plaque size compared to the POFUT2-HA tagged line reveals no difference between lines. Aiii) Numerical assessment of plaque capacity compared to the POFUT2-HA tagged line reveals no difference between lines. B) IFA assessment of localisation MIC2, detected using *α*-MIC2, and M2AP, detected using *α*-Flag, suggests that *O*-glycosylation plays no detectable role in protein trafficking. C) Invasion capacity assays at 2 and 10 minutes demonstrate no difference in invasion capacity.

Post-translational modifications can affect protein trafficking and we therefore assessed localisation of M2AP and MIC2 in *Δtgpofut2* by IFA. Here we could observe no difference in localisation of either MIC2 or M2AP suggesting that *O*-glycosylation plays no detectable role in protein trafficking (Figure 6B). We also specifically monitored for defects in host cell invasion after 2 and 10 minutes incubation on HFFs (Figure 6C). Again, we could see no difference in invasion capacity at either time point, using this assay, suggesting that TgPOFUT2 is not important for MIC2 function *in vitro*.

## Discussion

The recent recognition of the importance of glycosylation at the host-pathogen interface in parasites such as *P. falciparum* (19) and *Trypanosoma brucei* (29) has demonstrated the need to better understand glycosylation within phylogenetically distinct eukaryotic parasites. Here, we demonstrate that *T. gondii* has a functional POFUT2 homologue that is responsible for initiating *O*-glycosylation of TSR domains on MIC2. We confirm that at least three sites in MIC2 (S^285^, S^485^ and T^546^, Figure 3) are modified by GluFuc in addition to seven sites of *C-*glycosylation. These *O*-glycosylation sites lie within the previous proposed POFUT2 consensus sequon CXX(S/T)C (11,30,31) and suggest that TgPOFUT2 has a similar substrate preference to metazoan POFUT2 enzymes. Our data demonstrates that these sites are modified to a high occupancy with few peptide species lacking the *O*-glycans observed in parental MIC2 preparations. The loss of *O*-glycans in the *Δtgpofut2* line did not appear to impact *C-* glycosylation, as previously observed sites were readily detected within *Δtgpofut2* suggesting that in *T. gondii* MIC2 these two modifications operate independently of each other.

Previous work on mammalian POFUT2 has revealed that *O*-glycosylation is essential for mice with homozygous disruption of POFUT2 being embryonic lethal after implantation (15). In humans, loss of *O*-glycan elongation due to mutation in the β-1,3-glucosyltransferase (B3GLCT) (32,33) leads to Peter-Plus syndrome (34). These defects are driven by the requirement for the complete GlcFuc disaccharide for ER quality control where this glycan influences folding and stability of proteins with TSR domains (35,36). Similarly, we have observed that loss of *O*-fucosylation in *P. falciparum* sporozoites leads to the destabilization of the key adhesin TRAP (19,37). However, within *T. gondii* the loss of TgPOFUT2 lead only to a small change in MIC2 levels and no detectable difference in the complexing between MIC2 and its accessory protein M2AP or its localization. The small change in MIC2 levels is accompanied by no observable reduction in plaque size, which is consistent and the recent CRISPR-based fitness score assigned by Sidik *et al.* to TgPOFUT2 (−0.34 log_2_) (38), which in comparison to loss of MIC2 (−1.17 log_2_) is very mild. However, we cannot discount that TgPOFUT2 may be more important for *in vivo* (mouse) infection models where optimal tissue dissemination and invasion are critical for parasite survival. It may also be that TgPOFUT2 plays an important role in other stages of the parasite’s lifecycle (e.g. bradyzoites and enteric feline stages) which have not been assessed in this study.

Consistent with the non-essential nature of TgPOFUT2 in tachyzoites, we observed few changes across the proteome (Figure 4). These changes we observed were modest in magnitude, but consistent across replicates, suggesting that TgPOFUT2 targets a limited repertoire of substrates, in line with other POFUT2 enzymes [28, 33]. Interestingly, the absence of any dramatic decreases in abundance suggests that few proteins undergo destabilization as was observed with TRAP in *P. falciparum* [26]. This said, a number of proteins were observed to decreased within *Δtgpofut2* which may contribute to the observed phenotypes including TGGT1_270700, now known as *MYR2*, which was recently shown to be essential for effector translocation from dense granules [44]. We have not assessed effector translocation in this study and therefore this is worthy of more attention in the future. Additional we note that multiple members of the glycosylphosphatidylinositol-anchored SRS Superfamily [45] are altered in response to *Δtgpofut2* with an observed decrease in TGGT1_292270 (SRS36C) and an increase in TGGT1_292280 (SRS36D). As these surface antigens have been shown to modulate multiple aspects of infection [46] the function of these proteins may be worth following up. Surprisingly the most profound change we observed within the proteome of *Δtgpofut2* was an increase in the hypothetical protein TGGT1_237195. As this protein is not associated with *T. gondii* parasite fitness *in vitro* [42] and has yet characterized how or why the loss of *Δtgpofut2* results in an increase in abundance remains unknown. It is important to note that none of these proteins contain the TgPOFUT2 CXX(S/T)C sequon suggesting that they are not direct targets of TgPOFUT2 but may be being influenced indirectly by the loss of *O*-fucosylation. We did not see any difference in MIC2 levels in our global proteome analysis suggesting that this technique may not be sensitive enough to detect milder changes in protein levels.

The fact that TSR *O*-glycosylation does not appear to be important for the stability or trafficking of MIC2, in contrast to observations from TRAP in *P. falciparum* (19), may be due to the intimate association of MIC2 and M2AP. M2AP contributes to trafficking of this important adhesin to the micronemes and assists its function (39,40). Other coccidian parasites including *Neospora caninum* and *Eimeria tenella* also express orthologs of *T. gondii* M2AP (41,42). However, TRAP and related proteins in *P. falciparum* are not known to associate with an analogous protein and therefore it is plausible that this makes *O*-glycosylation more important for protein folding and trafficking in these species. We have also show here that MIC2 is heavily *C-* mannosylated, which alludes to the possibility that this modification could be important for the function of MIC2. This possibility is reflected by the fitness score assigned by Sidik *et al.* to the putative *C-*mannosyltransferase in *T. gondii* (TGGT1_280400) (−2.37 log_2_), which is suggestive of severe defects in tachyzoite function, though this remains to be investigated further.

In summary, TgPOFUT2 is responsible for *O*-glycosylation of TSR domains in MIC2. The loss of this modification leads to only small changes in MIC2 abundance, in contrast to the fate of TRAP in *P. falciparum* (19), and little to no impact on parasite virulence *in vitro*. Loss of TgPOFUT2 also provides no profound changes in the tachyzoite proteome. Taken together this demonstrates that TgPOFUT2 is dispensable for replication of *T. gondii* tachyzoites.

## Methods

### Plasmid construction and transfection

The 3’ portion of the *tgm2ap* (TGGT1_214940) ORF was PCR amplified using the following primers: 5’-TACTTCCAATCCAATTTAATGCTGCTTGAGCCGTGACAACAGATTAC-3’ and 5’-TCCTCCACTTCCAATTTTAGCCGCCTCATCGTCACTCGGCAGACGGC-3’ and LIC cloned into vector pSF-TAP.LIC.DHFR-TS as previously described (43). The pM2AP.SF-TAP construct was linearized within the *tgm2ap* homology region with *Pfl*MI prior to transfection into RH∆ku80:HXGPRT tachyzoites (43). The 3’ portion of the *tgpofut2* (TGGT1_273550) ORF was PCR amplified using the following primers: 5’-TAGTAGATCTAGCGATTAGCAC-TTTTTGG-3’ and 5’-AGCCCTAGGCAG-TGTCGAACTGGGGTC-3’ and ligated into the *Bgl*II/*Avr*II sites of pgCH (Gra 1 5’-CAT-*Bgl*II/*Avr*II 3xHA). The pgCH-POFUT2-3xHA construct was linearized within the tgpofut2 ho-mology region with *Mfe*I prior to transfection into RH∆ku80:HXG tachyzoites (43). This construct was linearised with Mfe1 (NEB) prior to transfection.

*Tgpofut2* was knocked out using a unique guide selected by EuPaGDT (http://grna.ctegd.uga.edu/batch_tagging.html) combined with a homologous repair template containing the BLE cassette. The CRISPR target plasmid (44) was constructed by Q5 mutagenesis (NEB) with the common reverse primer 5’-AACTTGACATCCCCATTTAC-3’ (45) and the forward primer 5’-GAGACGGTAAGAACTGAGACGGTTTTAG AGCTAGAAATAGCAAG-3’. The BLE cassette was then was then amplified using Primestar Max (Takara) containing homologous flanking regions to either side of the Cas9 cut site with primers 5’-TGTCTGCTCAACCACCGTCGCTTGCTGTTAGGCCTCGTCGTCTAGAAGGTGGATGCG GGA-3’ and 5’-TGAATGGGAGACACGAGAGGAAGACGGTAAGAACTGAGCGATGTGGAGTCGTCTC AAGCG-3’. 10 μg of the Cas9 plasmid was combined with 20 μg of PCR product, then precipitated using EtOH/NaAc prior to transfection. The dried DNA was resuspended in 3 μL of elution buffer (EB, Qiagen), followed by 20 μL P3 solution (Lonza). A washed parasite pellet containing approximately 10^6^ tachyzoites was then resuspended in this solution and transfected using the code FI-115 in a 16-well Nucleocuvette Strip in an Amaxa 4D Nucleofector (Lonza). Parasites containing the knockout construct were then selected as normal following addition of phleomycin and subsequently sub-cloned until a stable population was obtained. All transfections proceeded using either a Gene Pulser II (BioRad) or an Amaxa 4D Nucleofector (Lonza). Gene Pulser II transfection took place at 1.5kV and 25μF as is standard with 15µg of purified linearized DNA if seeking homologous integration.

### Parasite culture

Transfection and in vitro culture *T. gondii* tachyzoites were cultured under standard conditions. Briefly, Human Foreskin Fibroblasts (HFF, ATCC ^®^ SCRC-1041) were grown in DME supplemented with 10% heat inactivated Cosmic Calf Serum (Hyclone), until confluency was reached. Upon *T. gondii* infection HFFs with were refreshed with DME supplemented with 1% FCS. All cells were grown in humidified incubators at 37°C/10% CO_2_.

### Purification of M2AP-MIC2 complex

Parasites obtained from four T150 flasks confluent with HFFs were purified by filtration and collected by centrifugation (1000 g, 10 min). The parasite pellet was washed once on ice with PBS (2 ml). The parasite pellet was resuspended in 1 ml lysis buffer (50 mM Tris pH 8, 150 mM NaCl, 1% Triton X-100, Roche Mini Protease Inhibitor Cocktail, DNase) and lysed at room temperature for 20 min. The lysate was centrifuged (10000 g, 10 min, 4°C) to remove cellular debris and the supernatant incubated with Streptactin II resin (200 µl of 50% slurry) at 4°C for 1–2 h with nutation. The resin was collected in microspin columns (2000 g, 2 min, 4°C) and washed three times with 50 mM Tris pH 8, 150 mM NaCl, 0.1% Triton X-100. The resin was then incubated with 2.5 mM desthiobiotin, 50 mM Tris pH 8, 150 mM NaCl (200 µl per column, 20 min, 4°C) and the eluate collected by centrifugation (2000 g, 2 min, 4°C) to provide the purified MIC2-M2AP complex.

### Tryptic digest of gel-separated proteins

Affinity purified MIC2/M2AP was separated using SDS-PAGE, fixed and visualized with Coomassie G-250 according to protocol of Kang *et al.* (46). Bands of interest were excised and destained in a 50:50 solution of 50 mM NH4HCO3 / 100% ethanol for 20 minutes at room temperature with shaking at 750 rpm. Destained samples were then washed with 100% ethanol, vacuum-dried for 20 minutes and rehydrated in 50 mM NH_4_HCO_3_ plus 10 mM DTT. Reduction was carried out for 60 minutes at 56 °C with shaking. The reducing buffer was then removed and the gel bands washed twice in 100% ethanol for 10 minutes to remove residual DTT. Reduced ethanol washed samples were sequentially alkylated with 55 mM Iodoacetamide in 50 mM NH_4_HCO_3_ in the dark for 45 minutes at room temperature. Alkylated samples were then washed with two rounds of 100% ethanol and vacuum-dried. Alkylated samples were then rehydrated with 12 ng/µl trypsin (Promega) in 40 mM NH_4_HCO_3_ at 4°C for 1 hour. Excess trypsin was removed, gel pieces were covered in 40 mM NH_4_HCO_3_ and incubated overnight at 37 °C. Peptides were concentrated and desalted using C18 stage tips (47) before analysis by LC-MS.

### In-solution Glu-C/Trypsin digestion and double digestion

Affinity purified MIC2/M2AP was resuspend in 50ul 20% TFE and diluted equal volume of reduction/alkylation buffer (40mM TCEP, 80mM chloroacetamide and 100mM NH_4_HCO_3_). Samples were then heated at 40°C for 30 min to aid denaturation and reduction / alkylation in the dark. Glu-C or Tryspin was added (1/50 w/w) and allowed to incubate overnight at 37 °C. For double digestions after the initial Glu-C digestion trypsin (1/50 w/w) was added and allowed to incubate overnight at 37 °C. Digested samples were acidified to a final concentration of 0.5% formic acid and desalted using C18 stage tips (47) before analysis by LC-MS.

### Characterisation of MIC2 using Reversed phase LC-MS

Purified peptides prepared were re-suspend in Buffer A* and separated using a two-column chromatography set up composed of a PepMap100 C18 20 mm x 75 µm trap and a PepMap C18 500 mm x 75 µm analytical column (Thermo Fisher Scientific). Samples were concentrated onto the trap column at 5 µL/min for 5 minutes and infused into an Orbitrap Fusion™ Lumos™ Tribrid™ Mass Spectrometer (Thermo Fisher Scientific) or Orbitrap™ Q-exactive™ HF (Thermo Fisher Scientific) at 300 nl/minute via the analytical column using a Dionex Ultimate 3000 UPLC (Thermo Fisher Scientific). 75-minute gradients were run altering the buffer composition from 1% buffer B to 28% B over 45 minutes, then from 28% B to 40% B over 10 minutes, then from 40% B to 100% B over 2 minutes, the composition was held at 100% B for 3 minutes, and then dropped to 3% B over 5 minutes and held at 3% B for another 10 minutes. The Lumos™ Mass Spectrometer was operated in a data-dependent mode automatically switching between the acquisition of a single Orbitrap MS scan (120,000 resolution) every 3 seconds and MS-MS scan. For each ion selected for fragmentation Orbitrap HCD (maximum fill time 100 ms, AGC 2*10^5^ with a resolution of 30,000), Orbitrap EThcD (maximum fill time 100 ms, AGC 5*10^4^ with a resolution of 30,000) and ion trap CID (for each selected precursor (maximum fill time 100 ms and AGC 2*10^4^) was performed. The Q-exactive™ HF Mass Spectrometer was operated in a data-dependent mode automatically switching between the acquisition of a single Orbitrap MS scan (120,000 resolution) and 20 MS-MS scans (Orbitrap HCD, maximum fill time 100 ms and AGC 2*10^5^).

### Digestion of complex protein lysates for quantitative proteome

Parasites obtained from four T150 flasks confluent with HFFs were purified by filtration and collected by centrifugation (1000 g, 10 min). The parasite pellet was washed three times on ice with ice-cold PBS (2 ml). Parasites were lysed in ice-cold guanidinium chloride lysis buffer (6 M GdmCl, 100 mM Tris pH 8.5, 10 mM TCEP, 40 mM 2-Chloroacetamide) and boiled at 95 ˚C for 10 minutes with shaking at 2000 rpm to shear DNA and inactivate protease activity according to the protocol of Humphrey *et al.* (48). Lysates were then cooled for 10 minutes on ice then boiled again at 95 ˚C for 10 minutes with shaking at 2000 rpm. Lysates were cooled and protein concentration determined using a BCA assay. 100 ug of protein from each sample was acetone precipitated by mixing 4 volumes of ice-cold acetone with one volume of sample. Samples were precipitated overnight at −20 ˚C and then spun down at 4000 G for 10 minutes at 4 ˚C. The precipitated protein pellets were resuspended with 80% ice-cold acetone and precipitated for an additional 4 hours at −20 ˚C. Samples were spun down at 17000 G for 10 minutes at 4 ˚C to collect precipitated protein, the supernatant was discarded and excess acetone driven off at 65 ˚C for 5 minutes. Dried protein pellets were resuspended in 6 M urea, 2 M thiourea, 40 mM NH_4_HCO_3_ and reduced / alkylated prior to digestion with Lys-C (1/200 w/w) then trypsin (1/50 w/w) overnight as previously described (49). Digested samples were acidified to a final concentration of 0.5% formic acid and desalted using C18 stage tips (47) before analysis by LC-MS.

### Quantitative proteome of *Δtgpofut2* and parental lines using Reversed phase LC-MS

Purified peptides prepared were re-suspend in Buffer A* and separated using a two-column chromatography set up composed of a PepMap100 C18 20 mm x 75 µm trap and a PepMap C18 500 mm x 75 µm analytical column (Thermo Fisher Scientific). Samples were concentrated onto the trap column at 5 µL/min for 5 minutes and infused into an Orbitrap Fusion™ Lumos™ Tribrid™ Mass Spectrometer (Thermo Fisher Scientific) at 300 nl/minute via the analytical column using a Dionex Ultimate 3000 UPLC (Thermo Fisher Scientific). 180-minute gradients were run altering the buffer composition from 1% buffer B to 28% B over 150 minutes, then from 28% B to 40% B over 10 minutes, then from 40% B to 100% B over 2 minutes, the composition was held at 100% B for 3 minutes, and then dropped to 3% B over 5 minutes and held at 3% B for another 10 minutes. The Lumos™ Mass Spectrometer was operated in a data-dependent mode automatically switching between the acquisition of a single Orbitrap MS scan (120,000 resolution) every 3 seconds and MS-MS scans (Orbitrap HCD, maximum fill time 60 ms, AGC 2*10^5^ with a resolution of 15,000).

### Mass spectrometry data analysis

Identification of modification event within MIC2 and LFQ and was accomplished using MaxQuant (v1.5.3.1) (50). For the characterization of MIC2 searches were performed against the *T. gondii* (strain ATCC 50853 / GT1) proteome (Uniprot proteome id UP000005641, downloaded 02-04-2017, 8450 entries). For LFQ analysis searches were performed against the *T. gondii* (strain ATCC 50853 / GT1) proteome as well as the human (Uniprot proteome id UP000005640-*Homo sapiens*, downloaded 24/10/2013, 84,843 entries). For MIC2 searches carbamidomethylation of cysteine set as a fixed modification and the variable modifications of oxidation of methionine, *C-*glycosylation (+162.05 Da to W, allowing the loss of 120 Da due to the characteristic cross-ring fragmentation of the *C-*glycoside) and *O-*fucosylation (+308.11 Da to S or T, allowing a neutral loss of 308.11). For LFQ searches carbamidomethylation of cysteine was set as a fixed modification and the variable modifications of oxidation of methionine and acetylation of protein N-termini. Searches were performed with trypsin cleavage specificity allowing 2 miscleavage events with a maximum false discovery rate (FDR) of 1.0% set for protein and peptide identifications. To enhance the identification of peptides between samples the Match Between Runs option was enabled with a precursor match window set to 2 minutes and an alignment window of 10 minutes. For label-free quantitation, the MaxLFQ option within Maxquant (26) was enabled in addition to the re-quantification module. The resulting protein group output was processed within the Perseus (v1.4.0.6) (51) analysis environment to remove reverse matches and common protein contaminates prior. For LFQ comparisons missing values were imputed using Perseus and Pearson correlations visualized using R. All mass spectrometry proteomics data have been deposited to the ProteomeXchange Consortium via the PRIDE (52) partner repository with the dataset identifier PXD010714.

### Invasion Assay

Invasion assay proceeded as previously described (53). Briefly, Parasites were resuspended in a high [K^+^] buffer to supress motility, added to host cells allowed to settle. Buffer was then exchanged to DME supplemented with 1% FBS and parasites allowed to invade for 2mins and 10 mins at 37°C at which time were chemically fixed in 2.5% formaldehyde and 0.02% glutaraldehyde for 10 min each. The wells were then washed 3X with PBS pH 7.4 and were then blocked with 3% BSA in PBS pH 7.4 overnight at 4°C. IFA proceeded as standard using antibodies against SAG1 to detect extracellular parasites and then permeabilized with 0.1% Triton-X 100. GAP45 antibodies were then used to mark all parasites. Invasion rate was determined by counting the number of SAG1^+^ parasites as compared to total (GAP45^+^).

### Plaque Assay

Plaque assays were carried out by inoculating 120 parasites into each well of a 6-well plate in D1 media and allowed to grow undisturbed for 7-8 days. Cultures were fixed in 4% paraformaldehyde and incubated for 1hr at RT. Host cells were then stained using 2% crystal violet and washed.

## Acknowledgement

This work was supported by National Health and Medical Research Council of Australia (NHMRC) project grants awarded to NES (APP1100164). NES was supported by an Overseas (Biomedical) Fellowship (APP1037373) and a University of Melbourne Early Career Researcher Grant Scheme (Proposal number 603107). CJT was a recipient of an ARC Future Fellowship (FT120100164) that supported this work. We thank L. David Sibley for the kind gift of the pCas9-GFP plasmid and anti-MIC monoclonal antibody 6D10 as well as Giel Van Dooren for the kind gift of the pgCH plasmid. We declare no financial interests related to this work.

## Author contributions

E.D.G.-B and C.J.T conceived, designed all experiments and aided in the preparation of the manuscript. N.E.S performed all MS analysis, sample preparations, analyzed the results, and wrote the paper. M.J.C generated the *T. gondii* knockout lines. M.H.H and V.C generated M2AP-tagged line. A.J isolated MIC2. S.K and A.D.U preformed phenotypic assays.

**Supplementary Document: Supplementary Figure 1. Peptide coverage of MIC2 using our multiple enzymes approach.** Putative glycosylation sites are shown in red. Peptide containing *O-*fucosylated sites are highlighted in green.

**Supplementary Document: Supplementary Figure 2. Construct and validation of TgPOFUT2 within *T. gondii* GT1using cas9 driven mutagenesis.** Exon 1 of tgPOFUT2 was targeted for disruption using Cas9 driven mutagenesis. Introduction of a double strain break by guide enable the introduction of the BLE selectable marker

**Supplementary Document: Supplementary Figure 3. Pearson correlation of LFQ proteome experiments**. To assess the reproducibility of proteome analysis heat maps of observed LFQ proteome are provided. The mean Pearson correlation is >0.95

**Supplementary Document: Supplementary Table 1. Peptides observed from Trypsin, GluC and a combination of Trypsin followed by GluC digestion of parental MIC2.** MIC2 derived peptides from four digestions conditions revealed the presence of multiple glycosylation events, both *C* and *O-* glycosylation in MIC2. A total of four conditions were used to map glycosylation in MIC2; Tab 1) Trypsin reduced with DTT, Tab 1) Trypsin reduced with TCEP, Tab 2) GluC reduced with TCEP and Tab 3) Trypsin digestion followed by GluC digestion reduced with TCEP. For each condition the assigned Peptide sequence, Modification status, Mass, Protein and Maxquant assignment associated information (Score, Protein group, Scan number and data file name) are provided.

**Supplementary Document: Supplementary Table 2. Relative Quantitation of ^266^TLPQDAICSDWSA WSPCSVSCGDGSQIR^293^:** Multiple glycoforms of the peptide ^266^TLPQDAICSDWSA WSPCSVSCGDGSQIR^293^ are observable within MIC2. The most abundant of these forms all bear of *O-* fucosylation events support that S^285^ is modified at a high occupancy.

**Supplementary Document: Supplementary Table 3. Peptides observed from Trypsin of Δpft MIC2.** MIC2 derived peptides from a Tryptic digested reduced with TCEP. For this digest the assigned Peptide sequence, Modification status, Mass, Protein and Maxquant assignment associated information (Score, Protein group, Scan number and data file name) are provided.

**Supplementary Document: Supplementary Table 4. Relative Quantitation of ^266^TLPQDAICSDWSA WSPCSVSCGDGSQIR^293^ within Δpft strains:** Multiple glycoforms of the peptide ^266^TLPQDAICSDWSAWSPCSVSCGDGSQIR^293^ are observable within MIC2 yet no *O-*fucosylation were detectible.

**Supplementary Document: Supplementary Table 5. LFQ based analysis of ΔtgPOFUT2 vs parental strain.** A total of the 3839 protein groups observed across *T. gondii* proteome samples. For each protein group the observed LFQ values for all biological replicates, identification type, t-test significance, score, number of MS/MS events, iBAQ values and protein name gene generated using Maxquant are provided

